# 5’ Untranslated mRNA Regions Allow Bypass of Host Cell Translation Inhibition by *Legionella pneumophila*

**DOI:** 10.1101/2022.04.29.490120

**Authors:** Erion Lipo, Seblewongel Asrat, Wenwen Huo, Asaf Sol, Christopher S. Fraser, Ralph R. Isberg

## Abstract

*Legionella pneumophila* grows within membrane-bound vacuoles in alveolar macrophages during human disease. Pathogen manipulation of the host cell is driven by bacterial proteins translocated through a type IV secretion system (T4SS). Although host protein synthesis during infection is arrested by the action of several of these translocated effectors, translation of a subset of host proteins predicted to restrict the pathogen is maintained. To identify the spectrum of host proteins selectively synthesized after *L. pneumophila* challenge, macrophages infected with the pathogen were allowed to incorporate the amino acid analog azidohomoalanine (AHA) during a two-hour time window, and newly synthesized macrophage proteins were isolated by orthogonal chemistry followed by mass spectrometry. Among the proteins isolated were interferon-stimulated genes (ISGs) as well as proteins translated from highly abundant transcripts. Surprisingly, a large number of the identified proteins were from low abundance transcripts. These proteins were predicted to be among the most efficiently translated per unit transcript in the cell based on ribosome profiling datasets. To determine if high ribosome loading was a consequence of efficient translation initiation, the 5’ untranslated regions (5’UTR) of transcripts having the highest and lowest predicted loading levels were inserted upstream of a reporter, and translation efficiency was determined in response to *L. pneumophila* challenge. The efficiency of reporter expression largely correlated with predicted ribosome loading and lack of secondary structure. Therefore, determinants in the 5’UTR allow selected host cell transcripts to overcome a pathogen-driven translation blockade.

## Introduction

Diseases caused by intracellular pathogens remain significant and persistent global public health problems. Many of these organisms, such as *Mycobacterium tuberculosis* and the sexually transmitted *Chlamydia trachomatis*, grow intravacuolarly in membrane-bound compartments within host cells [1, 2]. *Legionella pneumophila* is one such Gram-negative intracellular pathogen that proliferates within host cells [3]. The organism was initially identified as the causative agent of Legionnaire’s disease, resulting from inhaled aerosols of water sources contaminated with amoebae bearing replicating *L. pneumophila*[4-6]. Once inhaled by the susceptible host, *Legionella* transfers its site of replication from amoebae to alveolar macrophages, with the potential for causing life-threatening atypical pneumonia [7, 8], particularly in immunocompromised individuals and the elderly[8].

The replication vacuole that bears *Legionella* avoids trafficking to the lysosome as a consequence of proteins translocated through the bacterial type IV secretion system [9-13]. Called Icm/Dot, this secretion system translocates over 300 proteins and is absolutely required for creating a replication niche and preventing host restriction[11, 14]. The translocated effector proteins promote the recruitment of endoplasmic reticulum (ER)-derived vacuolar components and modulate the host anti-pathogen response[15]. These include bacterial proteins that control host vesicular trafficking, block host cell translation and block the evolutionarily conserved host autophagy system[16].

The host can detect intracellular microbes through innate immune sensing by Pattern-Recognition Receptors (PRRs) such as Toll-like receptors (TLR) and Nod-like receptors (NLR), which recognize a variety of pathogen-associated molecular patterns (PAMP), including LPS and peptidoglycan derivatives[17]. Upon PRR activation, the innate immune response triggers the activation of transcription factors, such as those regulated by NF-κB, to transcribe pro-inflammatory cytokines and chemokines [18]. A common strategy used by intracellular pathogens is to inactivate components of the host protein synthesis machinery to undermine host restriction [10, 19-22]. Diphtheria toxin, shiga toxin, and *Pseudomonas* exotoxin A are examples of proteins that directly modulate host translation [23-25]. Although there are diverse explanations for the conservation of this tactic, it likely allows immune evasion when pathogens encounter hosts. *L. pneumophila* translocates at least seven proteins that depress host cell translation efficiency. Three of these proteins (Lgt1, Lgt2, Lgt3) are glucosyltransferases that modify and inactivate the host translation elongation factor 1A (eEF1a) [20]. As a consequence these secreted effector activities, a panel of host cell transcripts is induced that has been dubbed the *effector-triggered response* (ETR) [21]. This response involves the induction of both NF-κB as well as MAP kinase-dependent transcripts [26-31]. There also appears to be a host-driven response that inhibits translation initiation after *Legionella* infection that may have effects on transcriptional induction [32].

The inhibition of translation, which drives increased host cell transcription, has the paradoxical consequence that pathogen-response transcripts are predicted to be poorly translated due to elongation inhibition [28]. In spite of the low translation efficiency, pro-inflammatory cytokines such as tumor necrosis factor (TNF) and the interleukins IL-1α and IL-1β are clearly produced within cells harboring the bacterium [19, 28]. Arguing for the importance of this response in restricting bacterial growth, mice that are defective for IL-1α and IL-1β production and anti-TNF-α-treated rheumatoid arthritis patients are at high risk for *L. pneumophila* infection[19, 33]. These results are consistent with the model that the host is able to mount an immune response in the face of translation inhibition. Alternatively, the critical cytokine response necessary to clear the organism could be provided by uninfected bystander cells during disease, without the involvement of cytokines from infected cells, as has been shown in the mouse pneumonia model [34].

The ability of infected cells to produce inflammatory cytokines while the pathogen inhibits protein synthesis indicates that there must be mechanisms to allow translation in the face of these antagonists [28]. The primary accepted explanation for selective translation of inflammatory cytokines in infected cells is that the proteins synthesized in response to *L. pneumophila* originate from the most abundantly expressed transcripts in these cells [35]. This model argues that the synthesis of pro-inflammatory cytokines and chemokines occurs in infected cells as a consequence of 1000X transcriptional induction in response to *L. pneumophila*, allowing selective protein synthesis in the face of elongation inhibition[35]. Here we test this model by identifying proteins synthesized within the infected subpopulation of cells during a 2hr time period commencing at 4 hrs. post-infection (hpi). We show that in addition to proteins derived from highly transcribed genes, a large subset originates from genes having preferential ribosome loading without corresponding largescale transcription. As a consequence, efficient translation initiation bypasses the blockade caused by pathogen attack.

## Results

### Identification of host proteins that are selectively translated in the presence of *Legionella* infection

We identified the proteins encoded by mouse bone marrow-derived macrophages (BMDMs) that were selectively translated in response to *L. pneumophila*-GFP^+^ infection during a 2 hr. window. To this end, a snapshot proteomics strategy was performed, in which the methionine analog azidohomoalanine (AHA) was added from 4-6 hrs post-bacterial challenge [36]. The azido moiety on AHA provides a target for covalent linkage of newly synthesized proteins to alkyne-modified resin via orthogonal chemistry, permitting subsequent pelleting of resin and analysis [37] (Figure 1A). After the 2 hr incubation, BMDMs were harvested and sorted based on GFP fluorescence, a proxy for cells harboring *L. pneumophila*-GFP^+^. The BMDMs were lysed, and the lysate was subjected to orthogonal Cu^2+^ chemistry to allow linkage of newly synthesized proteins to the alkyne beads prior to pulldown and identification by LC-MS/MS (Materials and Methods). Prior to lysing, a portion of these cells were fixed, permeabilized, and AHA-labeled proteins were covalently linked to APC-phosphine to measure translation as a function of fluorescence intensity (Figure 1B). Infected cells, sorted on the basis of GFP fluorescence, showed a mean APC fluorescence intensity that was 5% the level of the bystander uninfected cells. Therefore, the sorted infected cell population that was subjected to proteomic analysis was strongly depressed for translation, consistent with previous observations on this population [28].

**Figure 1.**
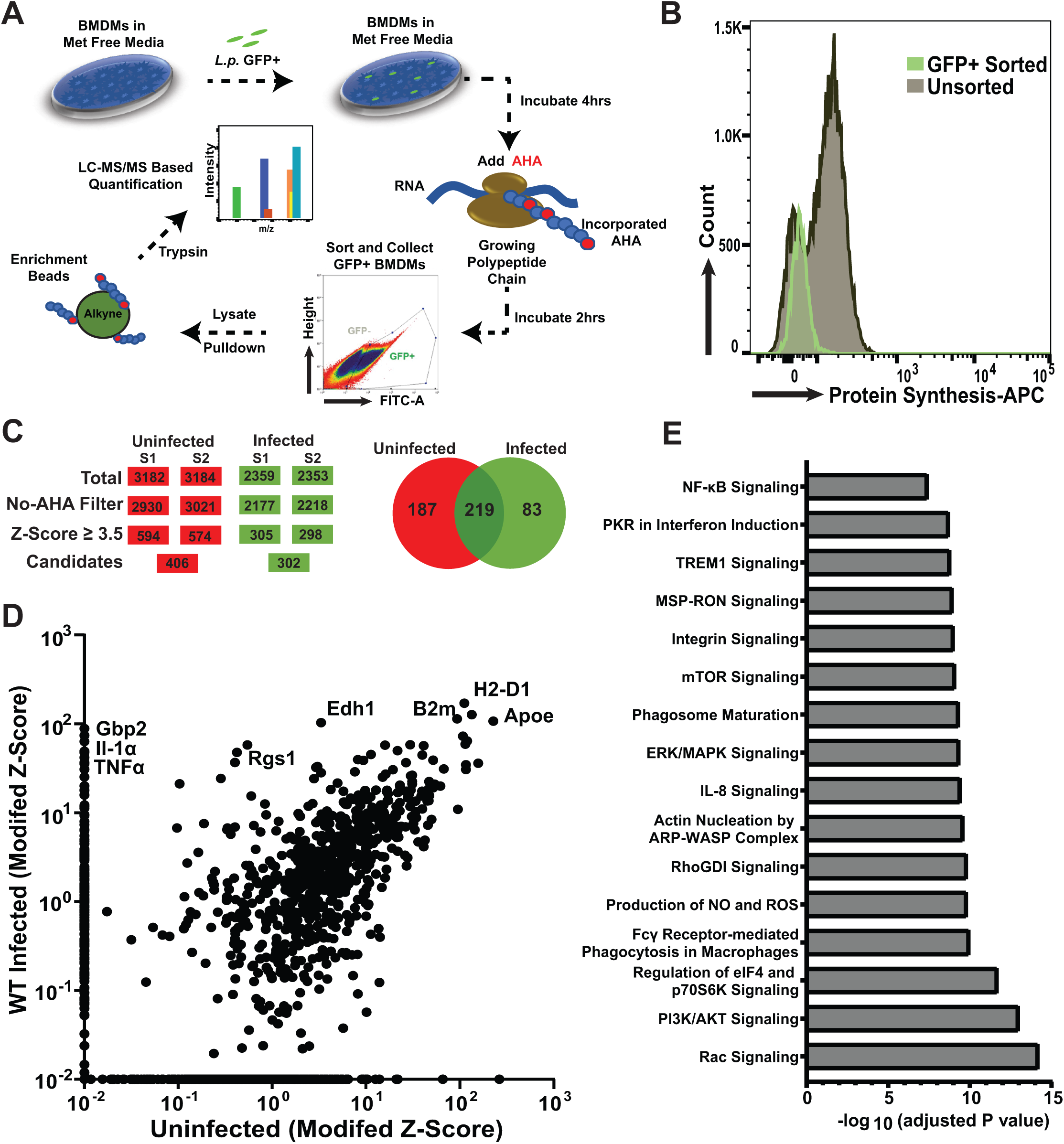
AHA labeling of BMDMs identifies 302 proteins selectively synthesized after *L. pneumophila* challenge. (A) Strategy for identification of newly synthesized proteins in macrophages infected with *L. pneumophila*. Methionine-starved C57Bl/6 BMDMs were challenged with *L. pneumophila-*GFP at a multiplicity of infection (MOI) of 15 and incubated for 4hrs. AHA was introduced into the medium, and infection was allowed to proceed for additional 2 hrs. Cells were then harvested and subjected to cell sorting, gating on GFP expression to identify macrophages with associated bacteria. The sorted cells from the GFP+ gate were lysed and incubated with alkyne-coated beads in the presence of Cu^2+^ to allow covalent linkage of AHA-incorporated nascent chains to resin (Materials and Methods). An uninfected sample was also run in parallel. The covalently linked proteins with AHA incorporated were released from the resin by trypsin digestion and then subjected to LC-MS/MS analysis to identify all proteins translated during this timeframe. (B) BMDMs challenged with *L. pneumophila* are blocked for protein synthesis. Shown is flow cytometry analysis of BMDMs cells infected with *L. pneumophila* at MOI = 15 for 6 hrs allowing for 2 hrs AHA incorporation as in panel (A). Sorted cells were fixed, permeabilized, and incubated with APC-phosphine to allow detection of AHA incorporation. (C) Flow chart demonstrating strategy for identification of proteins by mass spectrometry (MS). Modified Z-score analysis was performed (Materials and Methods)[72]. To determine nonspecific binding to the alkyne resin, a control sample was used with the omission of AHA. (D) Modified Z-scores for protein candidates identified by MS in uninfected and infected BMDMs. Proteins that were absent from a sample were given a value of 0.001 as the lowest limit of detection (Supplemental Dataset 1). (E) IPA-bases enriched pathway analysis of the list of newly synthesized proteins A (QIAGEN Inc, https://www.qiagenbioinformatics.com/products/ingenuitypathway-analysis) Proteins identified, values and statistical significance described in Supplemental Dataset 1.

Analysis of proteins proteolytically released from alkyne beads and analyzed by LC-MS/MS was performed on duplicate infections prepared on different days, with the rank-order results displayed as modified Z-scores for each candidate [38]. To identify candidates, a relatively stringent z >3.5 was used to ensure that extreme outliers will be retained as candidates, defined as proteins that show predominance relative to other components of the sample. To control for nonspecific binding to the alkyne resin, samples were used with the omission of AHA, and this group was filtered out from the outliers identified in the AHA-treated sample (Figure 1C). By comparing infected to naive uninfected BMDMs, we were able to identify 83 proteins that were overrepresented in cells harboring *L. pneumophila* (Figure 1D-E; Supplemental Dataset 1).

It should be noted that proteins identified by this method favor high molecular weight proteins and those that have a high representation of methionine residues. Despite this limitation, we were able to identify relatively small chemokines and cytokines in the GFP^+^ sample, such as TNF-α, that were not present in samples from cells not exposed to bacterium. Proteins identified by MS were submitted into various bioinformatics database searches [39-41]. This approach led to the determination that a large fraction of the candidates appeared to be interferon (IFN)-inducible proteins. The Interferome database contains IFN-stimulated genes (ISGs) curated from publicly available microarray datasets[42]. Roughly 60% of the proteins identified in the infected cells were IFN inducible based on these criteria (Figure 2C). This result is surprising because there is little or no IFN detectable by Western Blot or MS[43]; however, a number of reports have shown that there is low-level ISG expression as a consequence of basal IFN preexisting in cultured cells [43-45]. Focusing on the set of 83 proteins that was unique to infected BMDMs, 80% were ISGs.

**Figure 2.**
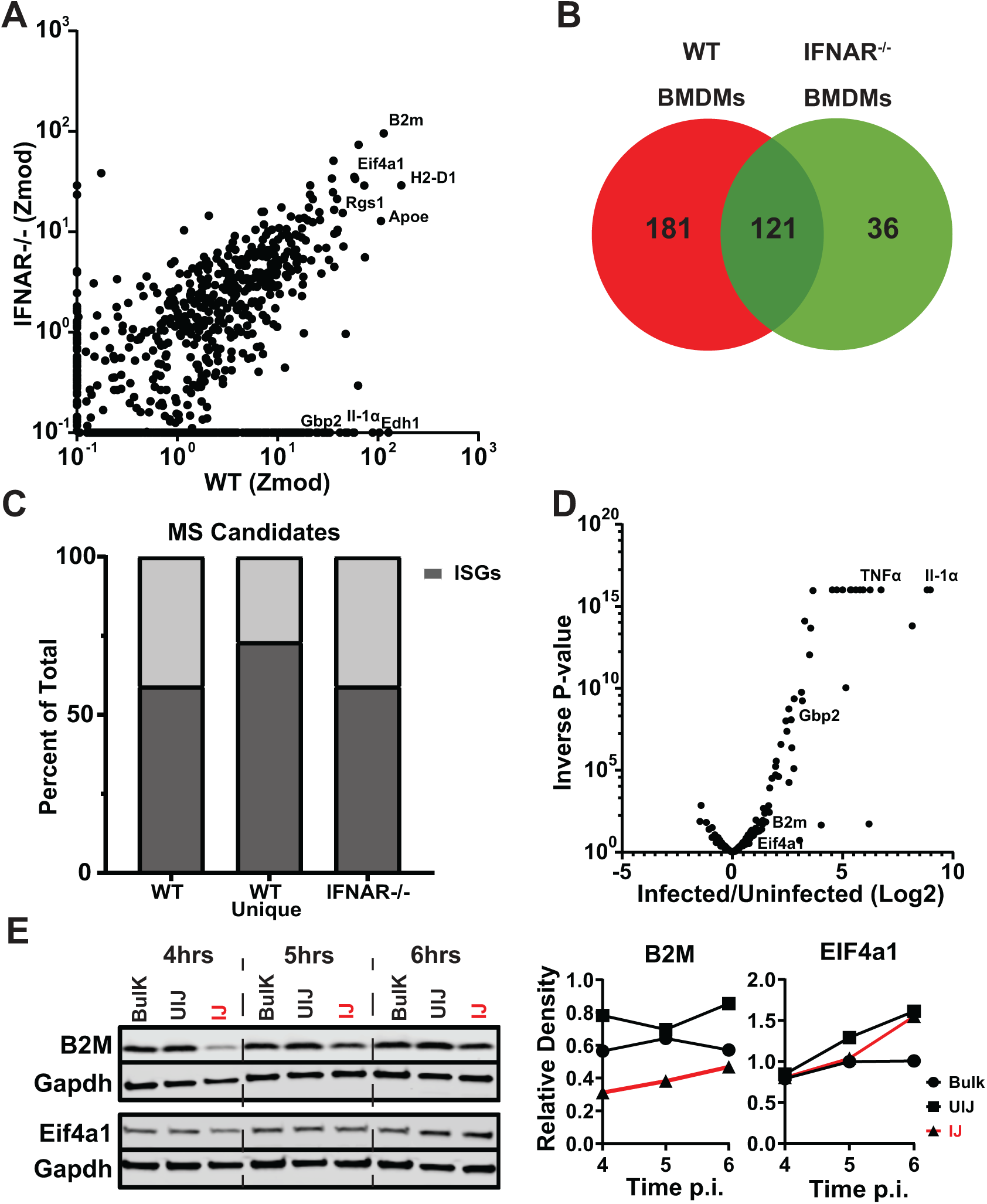
Dependence of identified translated host proteins on IFNa receptor signaling. (A) Modified Z-score analysis plot comparison between WT and IFNAR^-/-^ BMDMs infected with *L. pneumophila*. (B) Venn diagram comparing candidates with Z-score ≥ 3.5 in WT and IFNAR^-/-^ BMDMs challenged with *L. pneumophila*. (C) The ISG signatures of WT (total hits and hits unique to WT) and IFNAR^-/-^ BMDMs candidates as identified by the Interferome Database (Materials and ds). (D) Transcriptome analysis of BMDMs dataset[35] in the presence and absence of *L. pneumophila* infection. n is fold change in the presence of infection for the candidates identified by MS. 77% of candidates cluster (P>0.05, Test) with no evidence of transcriptional induction. Data for Figures 2A-2D detailed in Supplemental Dataset 2. (E) BMDMs were infected with *L. pneumophila*-GFP (Δ*flaA)* at MOI=15 and sorted 6 hours post-infection based on GFP ity. Immunoblot analysis of B2M and EIF4a1 both show no evidence of transcriptional induction after challenge. *pneumophila* WT. Bulk: total population before sorting; UI: uninfected (GFP-cells), IJ: infected (GFP+) cells. s on the right show densitometry in sorted cells normalized to Gapdh.

Given that the proteins were synthesized without obvious largescale IFN induction, the relative abundance of ISGs was further interrogated by challenging interferon-α/β receptor knockout (IFNAR-/-) BMDMs with *L. pneumophila*. IFNAR binds type I interferons and is necessary for upregulation of a set of ISGs in response to this cytokine. Proteomic analysis of IFNAR-/-macrophages was performed identically to the analysis of C57/BL6 (WT), labeling with AHA between 4-6 hrs post-infection, and the infected subpopulation of BMDMs was compared to that of IFNAR+/+ macrophages (Figure 2A; Supplemental Dataset 2). Overall, there was a 50% decrease in total proteins identified in IFNAR-/- BMDMs challenged with *L. pneumophila* compared to the WT (Figure 2 A-B). Surprisingly over 50% of proteins identified were still ISGs.

To understand if transcriptional induction of host genes upon *L. pneumophila* infection explained selective translation of these ISGs, RNA-sequencing data [35] in WT BMDMs was utilized to determine transcriptional differences amongst our candidates. Although a fraction of the candidates were transcriptionally upregulated, the majority (77%), however, show no statistically significant change in response to infection of BMDMs (Figure 2D). To verify that transcripts showing no increase in response to infection were able to continue translation in the presence of *L. pneumophila*, BMDMs were challenged with *L. pneumophila-*GFP^+^ and separated into infected (IJ) and bystander populations (UIJ). These samples were then probed at 4, 5, and 6 hrs post-infection with an antibody directed against B2M and EIF4a1 (Figure 2E), two proteins identified by MS to be encoded by transcripts that show no apparent induction in response to *L. pneumophila* challenge (Figure 2D). There was increased accumulation of both of these proteins over this time period after infection, indicating that these proteins were synthesized without corresponding transcriptional activation (Figure 2E; fraction IJ). Based on the result that a large subset of proteins originated from transcripts that were not transcriptionally induced, we next determined if those transcripts were simply expressed constitutively at high levels, or if a class exists that were translated from low abundance transcripts.

### One class of translated proteins is derived from abundant transcripts

Previous work noted that detection of immune response-related proteins in *L. pneumophila-*challenged BMDMs requires the presence of the pattern recognition receptor adaptor protein MyD88 [28, 46, 47]. Particularly striking was the fact that challenge of MyD88^-/-^ BMDMs results in high, although clearly attenuated, transcription of genes such as *Il1α* [28]. The dependence on MyD88 activity for detection of these proteins can be explained, as its absence abrogates the superinduction of transcripts, resulting in a correspondingly lower abundance of cytokine and immune response-related proteins[28, 35, 46, 48]. Although challenge of MyD88^-/-^ with *L. pneumophila* reduces the superinduction of transcripts associated with NFkB-regulated genes, it also allowed us an opportunity to investigate if there exists a class of proteins that are expressed in the presence of *L. pneumophila* without attendant superinduction.

BMDMs from C57Bl/6 (WT) and congenic MyD88^-/-^ macrophages were challenged with *L. pneumophila* and 6 hrs after infection, RNA was isolated and subjected to transcript analysis by mRNA-seq (Materials and Methods). Consistent with what has previously been reported [14, 49], challenge of WT BMDMs with *L. pneumophila* significantly induced transcription of genes encoding pro-inflammatory cytokines (*Tnf, Il1α, Il6*), chemokines (*Cxcl1, Cxcl2, Ccl3*), MAPK genes (*Dusp2, Dusp 8*), and NF-κB regulators (*IκB, Tnfaip3*) in WT macrophages (Fig. 3A, B; Supplemental Dataset 3). As expected, the overall induction of genes associated with the innate immune response was attenuated after *L. pneumophila* challenge of BMDMs from MyD88^-/-^ mice (Figure 3A, B). To support the model that transcript abundance allows bypass of translation inhibition, we measured absolute transcript levels in both WT and MyD88 deficient macrophages after challenge with *L. pneumophila* by calculating the number of aligned reads per kilobase of exon per million mapped reads (RPKM values) (Figure 3B). Transcription of pro-inflammatory cytokine (*Tnf* and *Il1*) and chemokine (*Cxcl2, Ccl2, Ccl4*) genes, which represented the most abundantly expressed genes in WT macrophages upon challenge with *L. pneumophila* (Figure 3B), were reduced 10-100x after challenge MyD88^-/-^ BMDMs, although the level of transcription of *Il1α* and *Il1β* remained relatively high compare to the total population [28, 50].

**Figure 3.**
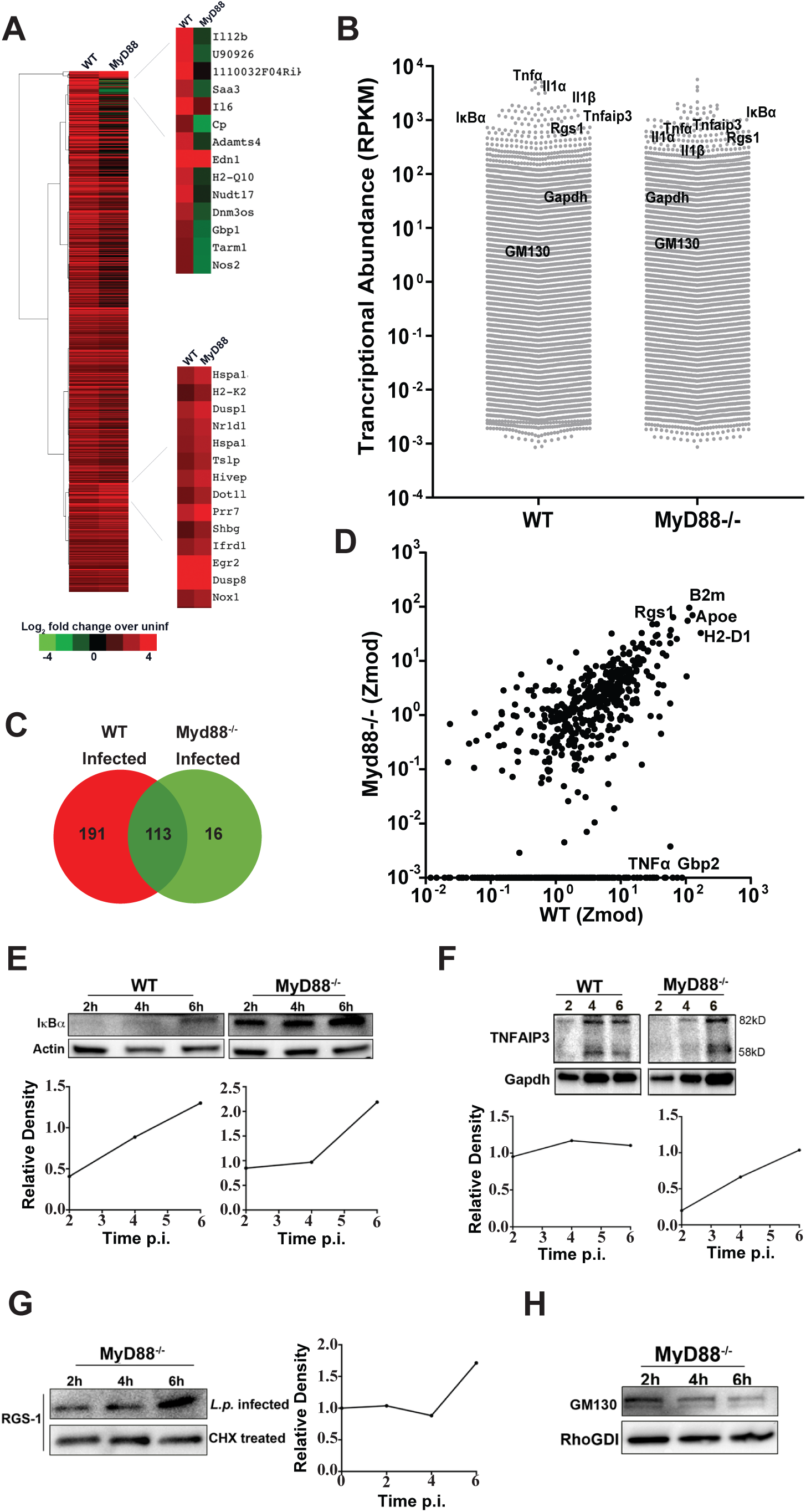
A subset of proteins encoded by highly abundant transcripts is translated in the presence of *L*.*pneumophila* infection. (A) Differential transcription in WT and MyD88^-/-^ macrophages infected with *L. pneumophila* Materials and Methods). Data are expressed as fold induction relative to samples of identical mouse macrophage strains incubated in the absence of bacteria. Left: Pearson hierarchical cluster analysis of 800 genes in WT and MyD88^-/-^ macrophages is displayed, showing genes with 2.5 fold change relative to control. Right: clusters showing either differential expression in WT vs. MyD88^-/-^ cells or genes similarly upregulated in response to infection in both cell types. (B) Absolute transcript abundance determined from Reads Per Kilobase of exon per million mapped reads (RPKM values) in macrophages harboring bacteria from the C57Bl/6 WT (WT) or MyD88^-/-^ harboring macrophages (MyD88). Each datapoint represents a single gene and a total of 22,006 genes were shown in each group. Genes are noted that show either high expression (in the top 5% of expressed genes) in WT infected BMDMs, with expression reduced 2-10 fold in Myd88^-/-^ BMDMs (TNFα, IL1α, IL1β, TNFAIP3, IκBα). In addition, genes with similar expression in both groups are noted (GM130, GAPDH). (C-D) Modified Z-score analysis plot comparison between WT and Myd88^-/-^ BMDMs infected with *L. pneumophila*. Protein candidates identified after infection with *L. pneumophila* with z-score ≥ 3.5 in WT and Myd88-/- BMDMs. (C) Venn diagram shows all candidates, not limited to those differentially transcribed. Figures 3A-D are from Supplemental Dataset 3. (E-H): WT or Myd88^-/-^ BMDMs were challenged with *L. pneumophila*-GFP (Δ*flaA*) at MOI=15 and sorted based on GFP^+^ 6 hours post-infection with flow cytometry. Immunoblot analysis of (E) IκBα and (F) TNFAIP3 in WT and MyD88^-/-^ infected and sorted cells. Bottom graphs show densitometry in sorted cells normalized to Actin and Gapdh, respectively. Immunoblot of RGS-1 (G) and GM130 (H) protein levels in MyD88^-/-^ macrophages challenged with *L. pneumophila* and sorted 2, 4, and 6 hrs after challenge (top lane). In parallel, cells were treated with 1 µg/mL cycloheximide (CHX) (G) (bottom lane). The graph on the right of panel G shows densitometry of RGS-1 in infected macrophages normalized to CHX treated cells.

To determine if transcriptional abundance is the sole determinant that drives bypass of the *L. pneumophila* translation block, Myd88^-/-^ BMDMs were challenged in duplicate with *L. pneumophila*, and the proteins translated between 4-6 hr post infection were identified by AHA incorporation. To identify the most abundant proteins, a rank order modified Z score ≥ 3.5 was used in the MyD88^-/-^ BMDMs relative to the WT. In total, there were 129 proteins identified in this fashion after Myd88^-/-^ BMDM infection, 90% of which were also found after infection of the WT (Figure 3 C, D). Furthermore, the 10% that did not meet the stringent significance cutoff after infection WT BMDMs could still be identified by MS in this infection, arguing against any unique proteins expressed in MyD88^-/-^ BMDMs. In contrast, about 60% of the proteins identified in the infected WT BMDMs had a modified Z-score < 3.5 from the MyD88^-/-^ sample. In fact, 25% of the proteins identified in the infected WT BMDMs were completely absent from MyD88^-/-^ sample (Figure 3D).

To demonstrate that the set of proteins identified after MyD88^-/-^ infection by MS continue to be translated during the course of *L. pneumophila* challenge, BMDMs were challenged with *L. pneumophila-*GFP^+^ for 2, 4, and 6 hours, with cells collected at each time point and sorted for the GFP^+^ infected populations. Extracts from the sorted cells were then immunoprobed with an antibody directed against IκBα (Figure 3E) and TNFAIP3 (Figure 3E-F), two proteins associated with negative regulation of the NFκB response that were abundantly transcribed in WT BMDMs after *L. pneumophila* challenge. In both WT and MyD88^-/-^ BMDMs, there was continued accumulation and increased steady-state levels of IκBα in spite of the presence of *L. pneumophila* protein synthesis inhibition (Fig. 3E) with similar results obtained with TNFAIP3 (Fig. 3F). RGS-1, the regulator of G-protein signaling-1 that is encoded by one of the most abundant transcripts in the MyD88^-/-^ BMDMs, showed similar levels of accumulation (Fig. 3G). To determine the amount of RGS-1 detected by immunoblotting that was due to *de novo* protein synthesis as opposed to long-lived species translated before infection, protein levels after infection were normalized to the amount of protein that remained during cycloheximide (CHX) treatment and plotted over time. Interestingly, significant accumulation occurred from 4-6 hrs post-infection, a time period previously shown to have maximum levels of protein synthesis inhibition [28]. In contrast to these examples, GM-130, which is transcribed at a level that is slightly above the median for MyD88^-/-^ BMDMs infected with *L. pneumophila*, showed no accumulation after infection (Fig. 3H). These results are consistent with the previously proposed model that transcript abundance in at least a subset of proteins is a determinant for bypassing translation inhibition during *Legionella* infection [35]. The results are also consistent with detection of proteins in this subset being dependent on transcription being above a minimum threshold.

### Identification of a set of proteins synthesized during *L. pneumophila* infection that are translated from poorly transcribed genes

To determine if a class of proteins exists that are synthesized efficiently without originating from abundant transcripts, we analyzed an extensive matched dataset that characterizes specific transcript abundancy by both RNA-seq and ribosome profiling after BMDM challenge with *L. pneumophila* [35]. We first compared the entire transcript pool to transcripts encoding the proteins that were identified during the 4-6 hr post-infection translation snapshot (Figure 4A; Supplemental Dataset 4). To analyze these datasets, the log10 transcript abundancy for these two populations were placed into bins, and the number of mapped transcripts was plotted as a function of their relative abundancy. There was a striking skewing of the transcript set identified by AHA labeling to the most abundant transcripts (Figure 4A). In fact, 70% of proteins identified in the translation snapshot were encoded by the most abundant 10% of transcripts. Furthermore, the top 2% of the most abundant transcripts contained 15% of MS-identified candidates. This analysis is consistent with the results arguing that transcript abundance is an important determinant for bypassing *L. pneumophila* translation inhibition [35].

**Figure 4:**
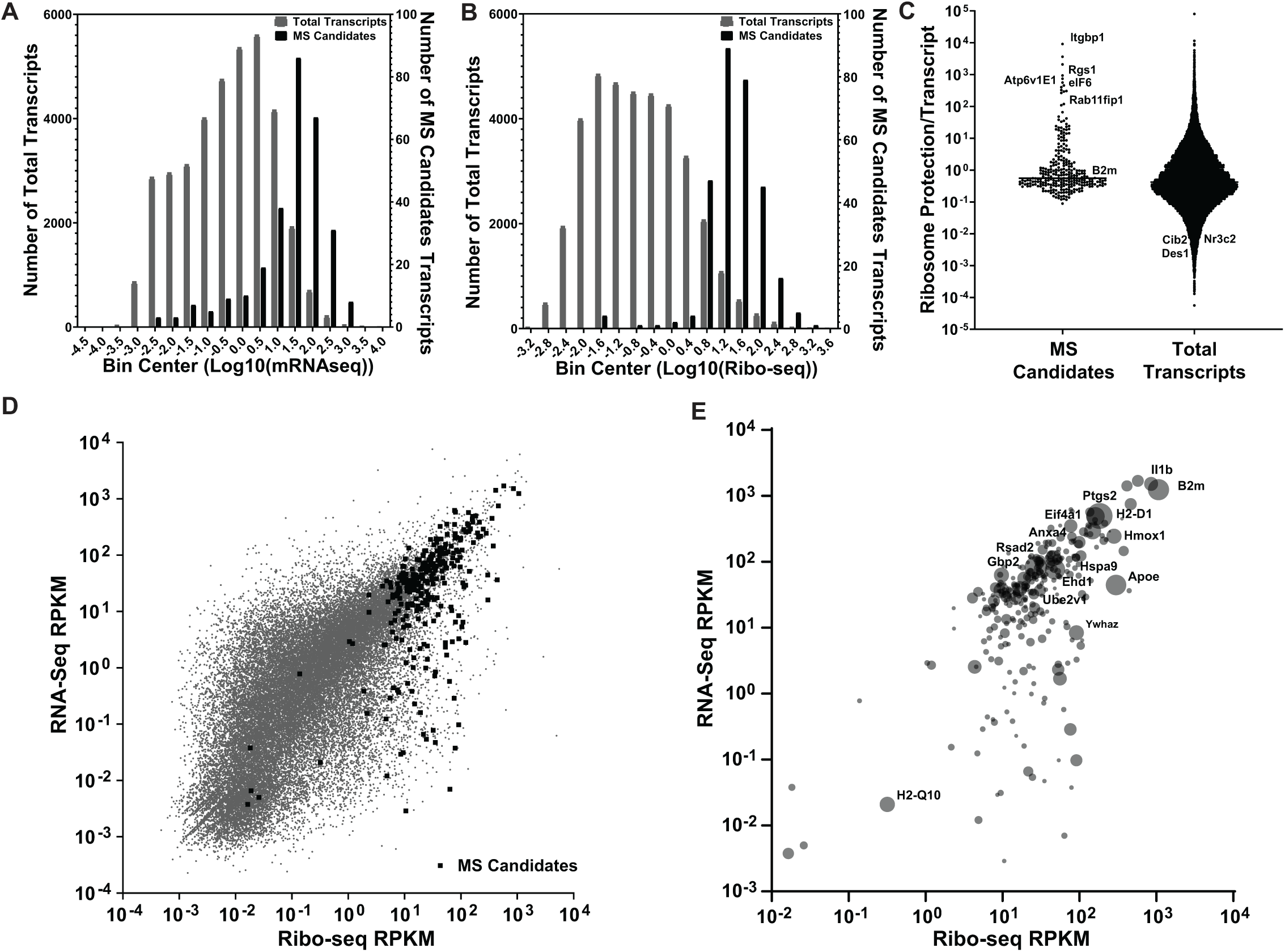
A class of proteins synthesized from low abundance transcripts is translated in the presence of *L. pneumophila* infection. (A-B) The mRNA-seq (RPKM) and ribosome profiling (RPKM) data are displayed for total transcripts (grey bars) [35] or proteins identified by mass spectrometry (Figs. 1,2; black bars). Number of total transcripts or MS candidate transcripts are displayed as a function of read density of transcripts (A) or ribosomal loading (B) [35]. Data on x axes are placed into 18 bins and based on processed raw data (Materials and Methods). (C) Translation efficiency (defined as (Ribo-seq RPKM)/(RNA-seq RPKM)) is displayed for each of the MS candidates and total transcripts. (D-E) Ribo-seq and RNA-seq read counts for total transcripts (gray circles) and for proteins identified by mass spectroscopy to be translated in the presence of *L. pneumophila* infection (black squares). (E) Data from panel D displayed as relative abundance of proteins identified by mass spectrometry. The size of the circle represents the protein abundance in the Mass-Spec. The top 15 most abundant hits have been labeled. RNA-seq and Ribo-seq data are in Supplemental Dataset 4.

Despite this general skewing, we still saw a considerable spread in the distribution of transcript abundance, with proteins that were translated from poorly transcribed genes found in the dataset. In addition, some proteins from highly abundant transcripts were missing after infection. For instance, IL-1α and TNFα were not observed in the MyD88^-/-^ samples even though their transcription levels were considerably higher than several proteins identified by MS (Figure 3B). In contrast, other proteins were encoded by some of the least abundant transcripts in the cell. Therefore, this dataset was investigated further to determine if there were other underlying molecular determinants that could explain the identification of proteins encoded by poorly transcribed genes. To this end, the available ribosome profiling (Ribo-Seq) [35] datasets from the same BMDM infections were used to determine if ribosome loading could be a determinant of bypass of *L. pneumophila* translation inhibition.

Using the identical strategy that we used to analyze transcription abundance, the number of transcripts in the cell was plotted as a function of the ribosome loading for that transcript from the published dataset [35], comparing the total pool of transcripts to those encoding the proteins identified in the 4-6 hr time window (Figure 4B; Supplemental Dataset 4). There was a powerful skewing of the proteins that we identified towards the most abundantly loaded transcripts. The top 5% of the most highly loaded transcripts contained 80% of MS-identified candidates. It should be noted, however, that the ribosome loading data is a count of the total amount of transcript protected from nuclease digestion by ribosomes, making the data a function of both the total amount of a particular transcript and the efficiency of loading on that transcript. In fact, when we measured loading efficiency per transcript, by expressing the data as the ratio of Ribo-seq protection/RNA-seq transcript abundance, the MS candidates from the WT infection showed a clear skewing toward heavier protection/transcript. In contrast, the entire population of transcripts showed a near normal distribution of protection (Figure 4C). Therefore, the MS candidates include proteins encoded by transcripts that show particularly efficient translation relative to the rest of the population.

Based on this analysis, there was a population of proteins identified in our MS dataset that appears to be more highly loaded than predicted from transcript abundance. To identify transcripts of low abundance that may be preferentially loaded, we displayed the relative transcript abundance as a function of the total ribosome loading and identified the transcripts that gave rise to the proteins identified after *L. pneumophila* infection (Figure 4D). The proteins identified that were encoded by poorly transcribed genes were predominantly skewed toward transcripts that were efficiently loaded. Therefore, these data argue that in addition to transcription abundance, there are sequence determinants that can allow poorly transcribed genes to be translated in the presence of *L. pneumophila* translation inhibitors.

The relative protein abundance determined by MS was next analyzed to determine if higher transcription levels were connected to yields determined by MS (Figure 4E). Although MS yields could have been affected by numerous confounding issues, such as efficiency of AHA incorporation as well as nonuniform fragmentation and peptide flight properties, there was general concordance between the highly expressed transcripts and protein abundance. Notably, proteins with high MS abundance were also identified that were modestly transcribed but heavily ribosome loaded (Fig. 4E). These included the 14-3-3ζ protein (YWHAZ) and apolipoprotein E (apoE), the latter of which was particularly abundant in the MS pool.

### Identification of 5’ UTR sequences that allow efficient translation of low abundance transcripts

Translation initiation is thought to be the rate-limiting step of protein synthesis, controlled by the efficiency of loading at the 5’ end of transcripts [51]. Given that a number of our identified candidates were predicted to be synthesized from heavily ribosome-loaded transcripts, we tested the model that the 5’ untranslated region (UTR) contributes to unexpectedly high abundance of proteins from a subset of poorly transcribed genes. We first searched for a consensus sequence in the 5’UTR of these top 20 loaded MS identify candidates but failed to identify such a sequence using a number of strategies (Materials and Methods). Surprisingly, we also did not observe a difference in 5’UTR length and GC content of these transcripts compared to poorly loaded transcripts (Supplemental Figure 1A-B). To determine if the 5’UTR contributes to translational efficiency, a series of reporter gene constructs were made that differed only in their 5’ UTRs. To this end, we set up a luciferase (Lux) reporter lacking a 5’UTR that is regulated by a 5X-NF-κB promoter (Fig. 5A) [27]. The 5’UTR from transcripts predicted to be translated either efficiently or inefficiently were then inserted upstream of the reporter, allowing the transcriptional fusions to be expressed specifically in response to *L. pneumophila* infection (pNL3.2.NF-κB-RE[NlucP/NF-κB-RE/Hygro]). In previous work, we demonstrated that the NF-κB promoter is quiescent in HEK293 cells in the absence of infection but becomes activated after exposure to *L. pneumophila*, dependent on the bacterial T4SS [27]. This allowed us to determine the amount of translation specifically after challenge with *L. pneumophila* without background contribution of translation prior to bacterial incubation. In a similar vein, as transcription from the reporter requires *L. pneumophila* incubation, quantitation of the *lux* transcript allowed accurate post-infection quantitation. This allows us to normalize the amount of protein synthesized to unit transcription.

**Figure 5:**
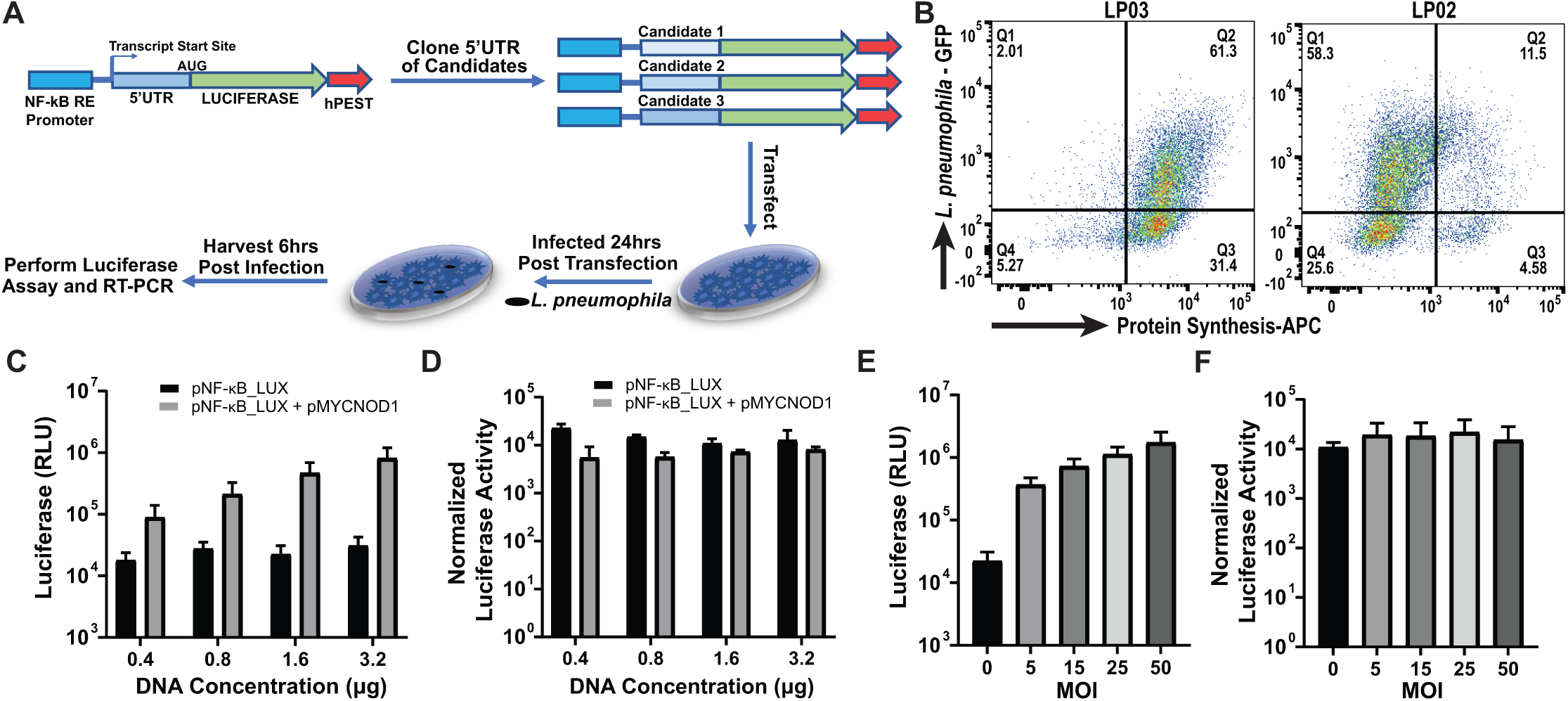
Strategy for determining translation efficiency as a function of 5’UTR sequence. (A) Experimental approach for evaluating the efficiency of translation driven by 5’UTR sequences after activation of an NF-kB responsive promoter in response to *L. pneumophila* challenge (B) Flow cytometric analysis of HEK-293T cells challenged with indicated *L* .*pneumophila* strains at MOI=15 for 4 hrs followed by 2 hrs further incubation in the presence of AHA. Cells were harvested, fixed, permeabilized, and labeled with APC-phosphine to detect and quantitate protein synthesis. (C-D) Luciferase activity after transfection with the indicated concentrations of the pNF-kB-LUX plasmid cotransfected with 800ng of pMYCNOD1. RT-PCR was performed to normalize data to the total mRNA level of luciferase and β-actin(D). (E) Luciferase activity after transfection with the 1.6 μg of pNF-kBLUX plasmid for 24 hrs and challenged with *L. pneumophila* at indicated MOI. RT-PCR was performed to normalize data to the total mRNA level of luciferase and β-Actin (F).

Individual 5’UTRs were selected from transcripts encoding candidates identified by AHA pulldown that were also predicted to have high levels of ribosome loading (Figure 4C), and regions starting at the 5’ nucleotide and ending precisely at the nucleotide preceding the ATG were inserted directly upstream of the *lux* reporter gene start codon. As controls, 5’UTR regions from transcripts shown to have low ribosomal loading were selected (Figure 4C). Each of the plasmids was transfected into HEK293-T cells, and after 24 hours, cells were either harvested for further processing or challenged with *L. pneumophila* for 6 hours. Luciferase activity was then measured as an indicator of mRNA translation, and the amount of *lux* transcript was determined by q-rtPCR. As we have argued previously [27], HEK293 behave very similarly to BMDMs with regard to the *L. pneumophila* translational block, as over 80% of the infected cells were shut down for translation, while a mutant lacking the T4SS (LP03) showed no translational interference (Figure 5B). These results were very similar to data from BMDMs [28].

The luciferase activity normalized to transcript concentration was stable over a range of DNA concentrations in the transfection mix, indicating that the normalized activity was independent of transcript levels (Figure 5C, D). The pNL3.2.NF-κB-RE[NlucP/NF-κB-RE/Hygro] with the native 5’UTR was cotransfected with increasing amounts of a plasmid encoding the Nod1 protein, a known inducer of the NF-κB promoter. Total luciferase activity was found to be dependent on DNA concentration (Figure 5C); however, when luciferase levels were normalized to transcript levels, there was no statistically significant differences between samples (Figure 5D). A similar test was performed by challenging transfected cells with increasing amounts of *L. pneumophila* (Fig. 5E, F). Total luciferase activity was found to be a function of MOI (Fig. 5E); however, when normalized to Lux mRNA, luciferase activity was independent of the increasing amount of transcription that occurred in these samples (Fig. 5F). Therefore, luciferase activity normalized to transcription level is predicted to be an accurate measure of translation efficiency.

To analyze the behavior of 5’UTRs, we chose B2M (one of the most abundant MS candidates) and five of the AHA-pulldown candidates that were translated from low abundance transcripts having high ribosome loading after *L. pneumophila* infection of C57/BL6 BMDMs (Figure 4C). As a control, three 5’UTRs from transcripts that resulted in low-efficiency loading were chosen (Figure 4C). The nine candidate 5’UTRs were fused upstream of Lux (Fig. 5A), transfected into HEK293 cells, and challenged with *L. pneumophila*, assaying for luciferase activity both before and after infection. Prior to infection, it was clear that most of the MS candidates showed higher basal activity than the controls (Figure 6A; underlying data, Supplemental Dataset 5). After 6 hr infection, five of the six 5’UTRs from transcripts encoding the MS candidates showed significantly higher normalized luciferase activities than the controls, which were defined as 5’UTRs from transcripts that showed low-efficiency loading based on ribosome profiling, with differences ranging from 2-10 times greater than the controls (Figure 6B, Supplemental Dataset 5; P = 0.05-0.001, depending on the reporter). The one exception was the 5’UTR of eIF6, which was among the MS candidates predicted to show high luciferase activity. The 5’UTR of eIF6 is unique amongst this subset as it contains an intron, so to determine if the presence of the intron interfered with our ability to see enhance translation, it was removed. However, removal of the intron did not result in increased normalized luciferase activity, indicating that the translation efficiency of the eIF6 transcript may be controlled by features outside of the 5’UTR. We conclude that the structure of the 5’UTR likely directs bypass of *L. pneumophila* translation inhibition in a large fraction of the transcripts encoding the MS candidates, but a subset of transcripts exist with other mRNA features that can facilitate bypass. Furthermore, the bypass is an intrinsic characteristic of the 5’ end of the transcript and not dependent on infection, as luciferase activities from transcripts encoding MS candidates were higher than the low-loading controls even in the absence of *L. pneumophila* challenge.

**Figure 6:**
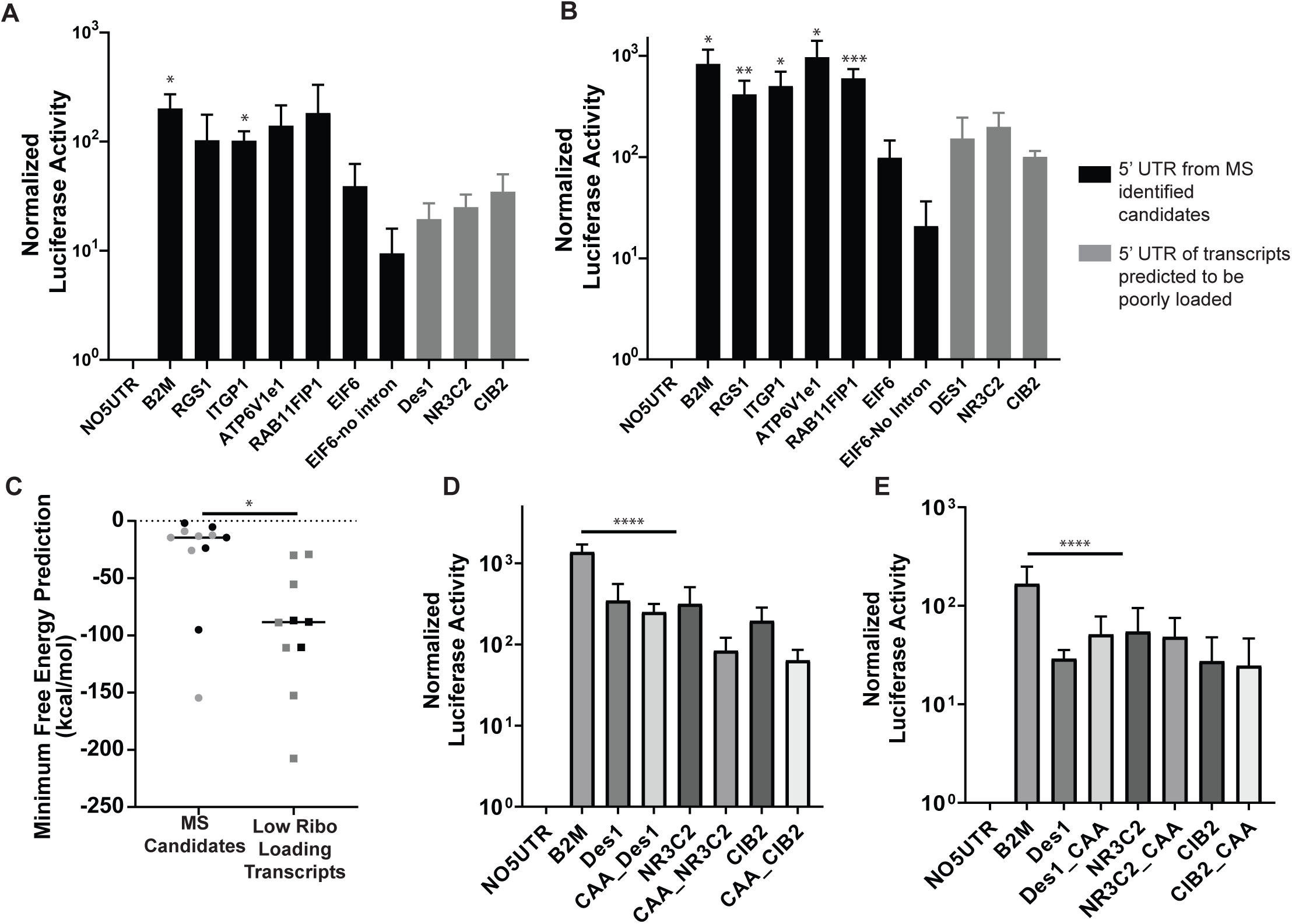
High ribosome loading transcripts have UTRs that enable efficient translation in the presence of *L. pneumophila* infection. (A) Luciferase activity after 24-hour transfection with 1.6μg of pNF-kBLUX plasmid containing selected 5’UTR transcript (n=3). qRT-PCR was performed to normalize data to the total mRNA levels of luciferase and β-Actin. Black bars: 5’ end from candidates identified by MS. Gray bars: 5’ end of transcript predicted to be poorly loaded [35]. (B) Transfectants having selected 5’UTR transcripts were challenged by *L. pneumophila* (MOI=25) and luciferase activity was determined after 6 hr (n=5). Data were normalized to total mRNA level of luciferase and β-Actin, as determined by qRT-PCR. Black bars: 5’ end from candidates identified by MS. Gray bars: 5’ end of transcript predicted to be poorly loaded. (C) Predicted minimum free energies for experimentally tested 5’UTRs (black) and 5’UTRs with similar ribosomal loading efficiency as the experimentally validated 5’UTRs (gray) (Materials and Methods)[75]. (D-E) 5’UTRs that drive low luciferase activity were modified with (CAA)_18_ at either the 5’ end of the transcript (D) or upstream of the translation start site, and constructs were transfected into HEK293T cells. The transfectants were then challenged by *L. pneumophila* for 6 hrs, and luciferase activities were determined (Materials and Methods). Statistical analyses were performed on normalized data by unpaired t-test (*< 0.05; **<0.01; ***< 0.001; Experimental Procedures).

As 5’UTR secondary structure has been proposed to play an essential role in regulating translational efficiency [52-54], we examined predicted secondary structures of each transcript variant that encodes the above experimentally validated 5’UTRs (Figure 6C, black points) and 5’UTRs with similar loading efficiency as the experimentally validated 5’UTRs (Figure 6C, gray points). Based on the Vienna RNA secondary structure algorithm, folding structures were predicted, as were the corresponding free energy changes [55]. The minimum free energy (MFE) for RNA folding of mRNA 5’UTR for this subset of genes was found to be significantly higher than poorly loaded transcripts, indicating that the poorly loaded transcripts likely have areas with extensive secondary structure not found in most of the efficiently translated transcripts (Figure 6C). These results are consistent with unstructured 5’UTRs conferring a translational advantage to transcripts in *L. pneumophila* infected cells.

As 5’UTR structures appear to regulate translational efficiency during *L. pneumophila* challenge of BMDMs, we manipulated the 5’UTR of the subset of transcripts that were poorly translated. Initiation factor eIF4F is recruited to the 5’ leader cap and collaborates with eIF4A to unwind the mRNA allowing for docking of the 40s ribosomal subunit[56]. Steric barriers in the 5’leader region can decrease translational efficiency, but it has been found that adding sequences lacking secondary structures either upstream or downstream of these structures, such as [CAA]_n_ repeats, can increase translational efficiency [57-60]. We inserted a [CAA]_18mer_ either proximal to the 5’UTR cap (Figure 6D; Supplemental Dataset 5) or upstream of the translation start site (Figure 6E; Supplemental Dataset 5). Insertion of these unstructured 5’UTR sequences was not sufficient to alleviate the translation suppression induced by the predicted secondary structures of these 5’UTRs during *L. pneumophila* challenge (Figure 6 C-D). Therefore, if secondary structure blockade interferes with ribosomal subunit docking, initiation of scanning by the preinitiation complex cannot overcome this blockage.

## Discussion

A number of pathogens encode proteins that interfere with host cell translation, leading to host cell misregulation, disruption of tissue organization and stimulation of innate immune signaling [22-25, 61]. The reasons for exerting translational control are varied, and range from supporting microbial colonization through localized inflammation to converting host translation into viral manufacturing platforms [62]. *L. pneumophila* is one such translation-blocking pathogen [21, 28, 63]. The mechanisms by which host cells can mount an immune response to restrict pathogen growth after translation blockade are poorly understood. In the case of host cellular responses to *Pseudomonas aeruginosa* and *L. pneumophila*, protein synthesis inhibition is tightly linked to transcriptional activation of antimicrobial signaling pathways, which provide the pool for residual translation in response to the pathogen. Recognition of microbial pattern molecules is central to this activation, as TLR signaling *via* MyD88 activates transcription, allowing bypass of protein synthesis inhibition [28, 35]. In addition, bystander cells, sensing either liberated microbial fragments or low-level cytokine production from infected cells, amplify this response and appear to be the primary source of inflammatory cytokine production in tissues [34].

In this work we determined if selective bypass of translational inhibition could be explained exclusively by transcriptional hyperinduction, resulting in secretion of innate immune effectors [28, 35]. To test this model, we identified the most abundant proteins synthesized from 4-6 hrs after *L. pneumophila* infection of macrophages, using snapshot proteomics analysis of the infected cell subpopulation. Among the over 300 identified proteins, a large number were synthesized from the most abundant macrophage transcripts (Fig. 4C), many of which required MyD88 for hyperinduction (Fig. 3C). This supports previous arguments regarding transcriptional hyperinduction [19], but does not fully explain *de novo* protein production. Proteins synthesized from transcripts that showed little or no induction in response to *L. pneumophila* were readily identified (Fig. 2D), and mutational inactivation of innate immune transcription driven by either Type I IFN (Fig. 2B) or MyD88 recognition (Fig 3C) did not prevent many of these proteins from being synthesized after bacterial infection. Most significantly, there was a group of proteins that were translated from poorly transcribed genes (Fig. 4D). Therefore, we believe that there are determinants other than transcriptional hyperactivation that contribute to bypass of *L. pneumophila*-dependent translation inhibition.

Our results provide evidence for translational control mechanisms that combat pathogen-directed inhibition of protein synthesis. Many of the proteins we identified after *L. pneumophila* challenge were derived from poorly-transcribed, highly ribosome-loaded transcripts (Figs. 4C, D) [19]. Surprisingly, although translational control of immune-related transcripts is potentially an important host strategy to combat pathogen attack, there has been minimal investigation of how the host uses ribosome loading to combat pathogen growth. Translation blockade could provide a significant hurdle for the pathogen, as viruses particularly are dependent on the host translation machinery for their replication[64]. Therefore, they must either overwhelm the host with transcripts, or evolve strategies to allow efficient translation of viral transcripts. It is logical to assume that the host has co-evolved mechanisms that similarly confer a translation advantage of immune-related transcripts in the face of translational attack. In particular, ISGs represent a first line of cellular defense against pathogens. Interestingly, we found that most of the proteins identified that were selectively synthesized after *L. pneumophila* attack are encoded by ISGs.

Preferential translation of a subset of transcripts can be explained by a number of molecular strategies that facilitate expression of immune response genes in the face of pathogen-encoded protein synthesis inhibitors. For example, it has been shown that translation efficiency could be altered by RNA structural elements such as upstream open reading frames (uORFs) or internal ribosome entry sites (IRES)[65]. The binding of translational regulatory proteins or non-coding RNAs within the 5’ end of transcripts also has the potential to modulate translation rates[66]. We argue, however, that in the presence of *L. pneumophila* challenge, that secondary structure of the 5’UTR is likely to be the important determinant that controls translation rates.

The frequency of uORFs or IRES elements appeared to be approximately the same in either efficiently loaded, poorly expressed or total cellular transcripts (Supplemental Figs 2, 3). In contrast, folding algorithms interrogating the 5’ untranslated regions (UTR) indicated that lack of secondary structure positively correlated with efficient translation (Fig. 6). This prediction was strongly supported by our analysis of 5’UTR fusions to a heterologous reporter, as MS candidates encoded by poorly transcribed genes had 5’UTRs that resulted in efficient luciferase production and lower secondary structure. The one exception to this rule, in which the 5’UTR of eIF6 was not sufficient to drive high luciferase production, argues that protein coding sequences can sometimes drive efficient protein synthesis in the presence of *Legionella* translation inhibitors. In fact, secondary structures located within the coding region have been shown to increase efficient loading of the ribosomal 43S preinitiation complex at the start codon, providing an alternate strategy that could explain efficient translation of eIF6 [57].

Previous work indicated that inhibition of translation by 5’UTR secondary structures can be reversed by increasing their distance from the 5’ end of the mRNA. This can be accomplished by inserting sequences lacking secondary structure, such as CAA repeats, between the 5’ end the element having high secondary structure [67]. The insertion of unstructured upstream regions is thought to facilitate scanning of the 5’UTR by the 43S preinitiation complex, allowing the AUG start codon to be identified [68]. Given that we found a strong correlation between unstructured 5’ UTRs and high translation levels, it is surprising that the addition of CAA repeats at the 5’ end of the poorly translated mRNAs failed to enhance translation efficiency (Fig. 6D-E). This result argues that the 5’UTR secondary structures found in poorly translated mRNAs may present profound blocks on scanning even under conditions in which scanning is efficiently initiated at the 5’ end. In support of this model, inhibitory secondary structure elements experimentally shown to be reversed by CAA repeats have calculated Δ*G* values of -30 to -50 kCal/mol [67], while most of the inefficiently translated 5’UTRs s in our study are predicted to have Δ*G* values that are considerably lower (Fig. 5). Alternatively, *L. pneumophila* infection could interfere with the function of translation initiation factors, preventing efficient movement through secondary structure elements after initial scanning of unstructured regions [32, 35].

In summary, this work provides evidence that efficient loading of low abundance transcripts is potentially an arm of the host defense against microbial infection. Many of these efficiently translated transcripts are interferon-regulated and/or fail to be induced in cells targeted by *L. pneumophila*. These results support previous work showing that low level transcription of antimicrobial factors is an important strategy for defending against pathogens that block translation or block innate immune response pathways in host cells[69]. Infected cells can then liberate either degraded microbial products or cytokines that signal to uninfected bystander cells to amplify the anti-pathogen response [34]. As pathogens are proficient at interfering with their recognition by host cells, this strategy is likely to be an important tool used by the host in the arms race with the pathogen.

## Materials and Methods

### Ethics Statement

This study was carried out in accordance with the recommendation in the Guide for Care and Use of Laboratory Animals of the National Institutes of Health. The Institutional Animal Care and Use Committee of Tufts University approved all animal procedures. The approved protocol number is B2013-18. The animal work, which is limited to macrophages’ isolation, does not involve any procedures of infections of live animals.

### Cell Culture

L cell supernatants were generated as described previously [70]. Bone marrow-derived macrophages (BMDMs) were from the femurs and tibias of female C57BL/6J as well as congenic MyD88^-/-^ and IFNAR^-/-^ mice (Jackson Laboratory, Bar Harbor, ME, USA). BMDMs were differentiated for seven days in Roswell Park Memorial Institute (RPMI, Gibco) medium containing 30% L cell supernatant, 10% Fetal Bovine Serum (FBS), 1mM L-glutamine, and 1x Penicillin/Streptomycin solution (100U/mL penicillin, 100µg/mL streptomycin) with feeding on day 4 of incubation. Cells were re-plated in antibiotic-free RPMI (10% L cell supernatant, 10% FBS, 1mM L-glutamine) medium 24 hours prior to infection with *L. pneumophila*. HEK293 cells (ATCC CRL-1573) were grown in Dulbecco modified Eagle medium (DMEM, Gibco) supplemented with 10% heat-inactivated fetal bovine serum (FBS).

### Bacterial Strains and Infection

All *L. pneumophila* strains are derived from LP02 [14], which is a streptomycin-resistant thymidine auxotroph. LPO2 Δ*flaA*-GFP+ (referred to as WT) and LPO3 Δ*flaA*-GFP+(referred to as *dotA3*) carry GFP on an isopropyl-β-D-thiogalactopyranoside (IPTG)–inducible, Cm-resistant plasmid (*ori*RSF101Cam^R^p*tac*::GFP^+^; referred to as pGFP). Strains containing the pGFP plasmid were maintained on BCYE plates [71]containing 100µg/mL thymidine and 5 µg/mL chloramphenicol and grown in AYE broth [71]containing 100µg/mL thymidine, 5µg/mL chloramphenicol, and 1mM IPTG.

For experiments involving challenge of cultured cells with bacteria, *L*.*pneumophila* colonies were patched on BCYE plates, and 36 hours later, twofold dilutions of *L*.*pneumophila* strains were grown overnight in AYE broth culture at 37ºC with shaking. Cultures were grown to post-exponential phase (A_600_ of 3.8-4.5), and dilution in this range was selected for challenge of mammalian cells. BMDMs were plated at a density of 1.56 × 10^5^ cells per cm^2^ and BMDMs were challenged at various multiplicities of infection (MOIs, assuming that A600 = 1.0 is equivalent to 10^9^ bacteria/mL. Contact was initiated by centrifugation for 10 mins at 400 rcf, and 1 hour post-infection, the medium was changed, and cells were maintained in the appropriate medium supplement with 200µg/mL of thymidine.

### Metabolic Labeling and Quantification

For all metabolic labeling experiments, BMDMs were plated at 1.56 × 10^5^ cells per cm^2^ on Costar Clear-Not-Treated 6-well plates (Corning, 3736) with RPMI containing 10% L cell-supernatant and 10% FBS. After 24 hours, the medium was changed to serum-free, methionine-free RPMI medium 1 hour prior to infection. Cells were challenged with *L. pneumophila* at MOI=15 by centrifugation for 10 mins at 400 rcf. After one hour, the cultured medium was changed to fresh RPMI (methionine and serum-free), and 50µM of L-azidohomoalanine (AHA) (Invitrogen) was added to medium at noted time points for a minimum of 1 hour incubation.

Cells were harvested at the indicated timepoints by placing on ice and washing twice with cold PBS. After the second wash, cells were harvested in cold PBS, pelleted, and fixed with 4% PFA for 15 mins. Fixed cells were washed twice with PBS and blocked with BSA/PBS for 1 hour.

After 1 hour, 100μM of APC-conjugated phosphine reagent (Pierce) in blocking buffer was added. Cells were incubated at 37ºC for 2-3hrs, and washed three times with 0.5% Tween-20/PBS followed by analysis on a Becton-Dickenson FACScalibur.

### Identification of newly synthesized proteins by click chemistry and mass spectrometry

To identify newly synthesized proteins in response to *L. pneumophila* challenge, BMDMs were plated at 1.56 × 10^5^ cells per cm^2^ on Costar Clear Not Treated 6-well plates (Corning, 3736) with RPMI containing 10% L cell supernatant and10% FBS. Cells were changed to serum-free, methionine-free RPMI medium 1 hour prior to infection, then challenged with *L. pneumophila* Δ*flaA*-GFP^+^ at MOI=15. The medium was then changed to fresh methionine-free RPMI 1-hour post-infection followed by addition of 100µM of L-azidohomoalanine (AHA) at 4 hours post-infection. After 2 hrs further incubation, cells (2.4×10^7^) were then placed on ice, lifted in 2ml cold PBS by pooling two 6-well plates and sorted by BD FACSAria. 1×10^7^ GFP^+^ cells were the collected, pelleted, and flash frozen for further proteomic analysis.

Pulldown of newly synthesized proteins using the Click-iT Protein Enrichment kit followed manufacturer’s protocols (Thermo-Fisher). The flash-frozen cells were lysed by resuspending in Urea lysis buffer (8M Urea, 200mM Tris=pH 8.0, 3% CHAPS and 1M NaCl, supplemented with protease inhibitors, and then treated with Benzonase (2.4U/mL) followed by incubation on ice for 15-30min. The lysate was then vortexed for 5 mins, debris pelleted at 10,000 RCF for 5 minutes, and the supernatant was transferred to a fresh tube for further analysis.

To set up the Click Reaction, 2x catalyst solution was mixed with the 800µl of cell lysate and 200µl of resin slurry. The sample was rotated end-over-end at room temperature for 20 hours and resin was pelleted by 1 minute centrifugation at 1,000 RCF, and the supernatant was removed. 1ml of SDS wash buffer (Click-iT Protein Enrichment kit) containing 10µl of 1M DTT was added to the resin and then heated to 70ºC for 15 mins prior to incubation at room temperature for 15 mins. The resin was pelleted for 5mins at 1000 RCF, and the supernatant was removed. The 1ml resin-bound protein fraction was then incubated with the addition of 7.4mg of iodoacetamide and incubated in the dark for 30 mins. The resin was pelleted, transferred to a column for washing. The column containing the beads was washed 5x with SDS Wash buffer (Click-iT Protein Enrichment kit), 5x with 8M Urea/100mM Tris pH=8, 5x with 20% acetonitrile in water. To digest the resin-bound protein, the resin was resuspended in 500μl of digestion buffer (100mM Tris-HCl (pH=8), 2mM CaCl_2_ 10% acetonitrile). Then 0.1 µg/µL of trypsin was added to the resin slurry, and the mixture was incubated at 37°C overnight. After the overnight digestion, the resin was pelleted by centrifugation for 5 minutes at 1000 RCF, the supernatant was collected, and the samples were acidified with 2µl of TFA to stop further reaction.

Samples were submitted to the Taplin Mass Spectrometry Facility (https://taplin.med.harvard.edu/home) for LC-MS/MS. To account for the mass gain due to incorporating the methionine analog, AHA sample parameters were modified to account for either a 107 AMU mass gain if the analog is incorporate instead of methionine or the incorporation of endogenous methionine. The LC-MS/MS was performed in duplicate, and the results were analyzed using modified Z-score to rank-order candidates. Other strategies, such as individual peptide abundance, gave similar results. To determine the *z*_MOD,_ the deviation of each protein from the median intensity of proteins in the sample was determined as the Median Absolute Deviation (MAD). From this calculation, the *z*_MOD_ was determined as described[72] and outliers were determined as *z*_MOD_ ≥ 3.5. Outliers were then interrogated using Ingenuity Pathway Analysis (IPA) database (QIAGEN Inc, https://www.qiagenbioinformatics.com/products/ingenuitypathway-analysis) which categorizes proteins based on function, pathway, and network. Candidates from the LC-MS/MS analysis were further dissected using Interferome (http://interferome.its.monash.edu.au/interferome/home.jspx, version 2.01).

### Immunoblot analysis of protein levels in infected BMDMs lysates

BMDMs were plated at 1.56 × 10^5^ cells per cm^2^ on Costar Clear Not Treated 6-well plates with RPMI containing 10% L cell supernatant/10% FBS. After 24 hours, cells were challenged with *L. pneumophila* at MOI-15, followed by contacting the BMDMs in the centrifuge at 400 RCF for 10 min. At the indicated timepoints, medium was removed and adherent cells were lifted with ice cold PBS and pelleted at 1000 RCF for 5 mins in the centrifuge. Cells were sorted and lysed with 1x SDS Laemmli sample buffer (0.125 M Tris-Cl pH 6.8, 4 % SDS, 20% glycerol, 10% beta-mercaptoethanol, 0.01 % bromophenol blue). Cell lysates were boiled for 5 mins and fractioned on SDS-polyacrylamide gels (Bio-Rad) and then electroblotted on nitrocellulose membranes. Blots were blocked with 5% BSA in Tris-Buffered Saline-Tween (TBST: 0.05 M Tris-HCl (pH=8.0), 0.138 M NaCl, 0.0027 M KCl, 0.05% Tween-20). For immune detection, cells were probed with a primary antibody (1:1000 or manufactures’ recommendation) in 5% BSA/TBST overnight at 4ºC. After washing with TBST, secondary antibodies Dylight anti-rabbit IgG 680, Dylight anti-mouse IgG 680, Dylight anti-rabbit IgG 800, or Dylight anti-mouse IgG 800 (Cell Signaling 1:20,000) were incubated in 4% milk/TBST for 45 minutes at room temperature. The membranes were scanned by Odyssey Imaging System and the Image Studio software (LI-COR Biosciences).

### Processing of Sequencing Data

BMDMs from WT and MyD88^-/-^ mice were challenged with LPO2 Δ*flaA*-GFP^+^ at MOI = 3 and cells harboring bacteria were sorted by flow cytometry based on GFP fluorescence (Fig. 3A-D). RNA was extracted from sorted cells, DNase treated (Turbo DNA-free kit, Invitrogen) and used for generating RNA-Seq library using TruSeq stranded total RNA library prep kit (Illumina). cDNA fragments from the library were sequenced by Illumina HiSeq 2000 (150bp, single end reads). RNA sequencing reads were processed using CLC Genomics Workbench (Qiagen). Reads were preprocessed by first trimming the linker sequence from the 3’end, and then aligned to the mouse genome (mm10). For read mapping parameters, a maximum of four mismatches were allowed, and multi mapping of up to eight different positions was permitted. mRNA transcription track alignment was performed. Only one genomic position per alignment was allowed. Reads per Kilobase of Transcript per Million mapped reads (RPKM) value was calculated as the expression value. RNA-seq data were deposited in BioSample (SAMN10180267, SAMN10180268). RNA-seq and Ribo-seq data for *L. pneumophila* challenged C57Bl/6 BMDMs were also obtained from a published dataset, GSE89184 (Fig. 4) [35].

### 5’UTR Sequence Analysis

5’UTR sequences of selected transcripts were obtained from Ensemble genome browser (http://useast.ensembl.org/index.html, Ensembl Genes 103, GRCm39) in FASTA format. In the cases in which more than one transcript was available, the transcripts with the highest ribosomal loading were chosen. The MEME algorithm present at the MEME suite database (University of Nevada, Reno, University of Washington, Seattle, WA, USA, Version 5.3.3; http://meme-suite.org/) was used for identification of sequence motifs in a collection of unaligned nucleotide sequences [70]. Possible uORF and IRES sequence features were analyzed by comparing them to a publicly available uORF-containing dataset [73]and IRES containing dataset [74]. GC content was calculated as a percentage-based formula: Count(G+C)/Count(A+T+C+G) * 100.

Secondary structure and minimum free energy was predicted by RNAfold (Institute for Theoretical Chemistry, University of Vienna, Vienna, Austria, http://rna.tbi.univie.ac.at/cgi-bin/RNAWebSuite/RNAfold.cgi) [75]

### Construction of Reporter Plasmids

A sequence spanning the transcription start site (TSS) to the translation start site of the selected 5’UTRs was amplified by PCR from mouse BMDM genomic DNA using the primers listed in Supplemental Table 1, containing flanking sequences that matched the pNL3.2NF-κB-RE plasmid (N111A, Promega). The pNL3.2NF-κB-RE plasmid was PCR amplified with primers containing sequences flanking the 5’UTR. Amplification was carried out in a PCR Thermal Cycler (Thermo Scientific) with a preliminary denaturation step at 94 °C for 5 min, followed by 30 cycles at 94 °C for 45s, primer annealing at 60°C for 15s and primer extension at 72°C for 30s, followed by a 2-min final extension at 72°C. PCR products were cleaned using QIAquick Gel Extraction Protocol (Qiagen). The PCR fragment and the vector were gel extracted and combined in a Gibson Assembly Reaction (NEB, E2611S) and transformed into DH5α. Clones were sequenced, and positive clones were stored in -80ºC.

### 5’UTR Luciferase Activity Reporter Measurements During *L*.*pneumophila* Challenge

HEK293 cells were plated at 1 × 10^5^ on 12 well plates in DMEM (10% FBS). Twenty-four hours after seeding, cells at ∼80% confluency were subject to transfection using Lipofectamine 2000 reagent (Invitrogen) according to the manufacturer’s protocol. The cells were either transfected with the indicated concentration of pNL3.2NF-κB-RE plasmid reporter construct alone or co-transfected with pMYCNOD1. Twenty-four hours post-transfection the medium was replaced, and cells were challenged *L*.*pneumophila* at the indicated MOI followed by 400 RCF centrifugation for 10 minutes. After 1-hour, the medium was replaced, and the cells were incubated for another 5 hours. Cells were harvested 6 hours post-infection with cold PBS. Two washes with PBS were performed, and the cells were resuspended in1ml of cold PBS. The sample was split into two aliquots; 100μl aliquots were used for luciferase measurements, 900µl aliquots were used for RNA extract and RT-qPCR.

Luciferase activity was quantified using the Nano-Glow Luciferase Assay System (Promega) according to the manufacturer’s instructions. 100μl aliquots were transferred to a Corning 96-well White Flat Bottom polystyrene plate for luciferase measurements.

Luminescence was measured using the Synergy Microplate reader (BioTek Instruments) and was determined as relative luminescence units (RLUs). Briefly, one volume of Nano-glow Luciferase Assay Reagent equal to the sample volume was added. The mixture was incubated for 3 minutes, and the luminescence intensity was measured. To correct for differences in transfection efficiency, luciferase activities were normalized to luciferase mRNA transcript values and β-actin transcript values in each sample.

For total RNA preparation, to determine luciferase activity, 900µl cell samples were pelleted and RNA was extracted from cells by using the RNeasy kit (Qiagen). The resulting total RNA sample was diluted to 1μg of total RNA in 10μl of H_2_O and treated with ezDNASE (Invitrogen) enzyme to digest gDNA. cDNAs were synthesized using SuperScript IV VILO Reverse Transcriptase kit (Life Technologies) with random primers using 1µg of RNA as a template. Each cDNA sample was used as a template to analyze luciferase transcript level using primers in Table 1. The expression level of luciferase was standardized by normalizing to the expression levels of β-Actin. SYBR Green PCR Master Mix reagent was used to perform Quantitative PCR.

## Acknowledgements

We would like to thank Mr. Ross Tomaino of the Taplin Mass Spectrometry Laboratories at Harvard Medical School for responsive proteomic analyses, and Drs. Russell Vance, Wenwen Huo, Philipp Aurass and Seongok Kim for review of the text. EL was supported by training grant T32-GM007310 from NIGMS. Work was supported by R01-AI146245 and R01-AI113211 from NIAID.

---

**Supplementary Figure 1.**
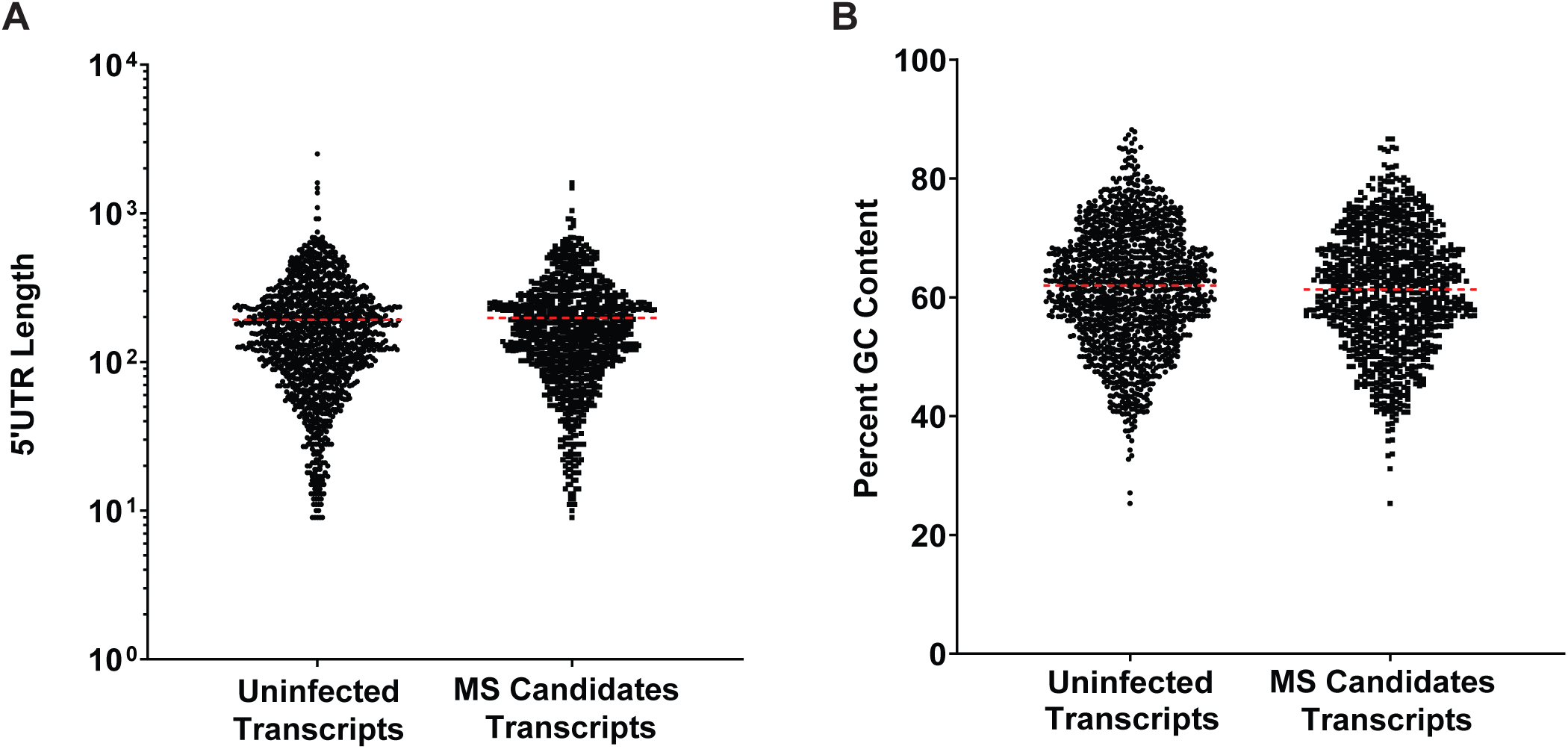
Distribution of Length and GC content of 5’UTR of MS candidates is unaltered relative to uninfected controls. (A-B) Comparison of the RefSeq-annotated 5’UTR lengths (A) and GC content (B) for transcripts of proteins identified by MS. The dot plots depict the distribution of 5’UTR length and GC content with no statistical significance between samples. Unpaired t-test statistical analyses were performed.

**Supplementary Figure 2.**
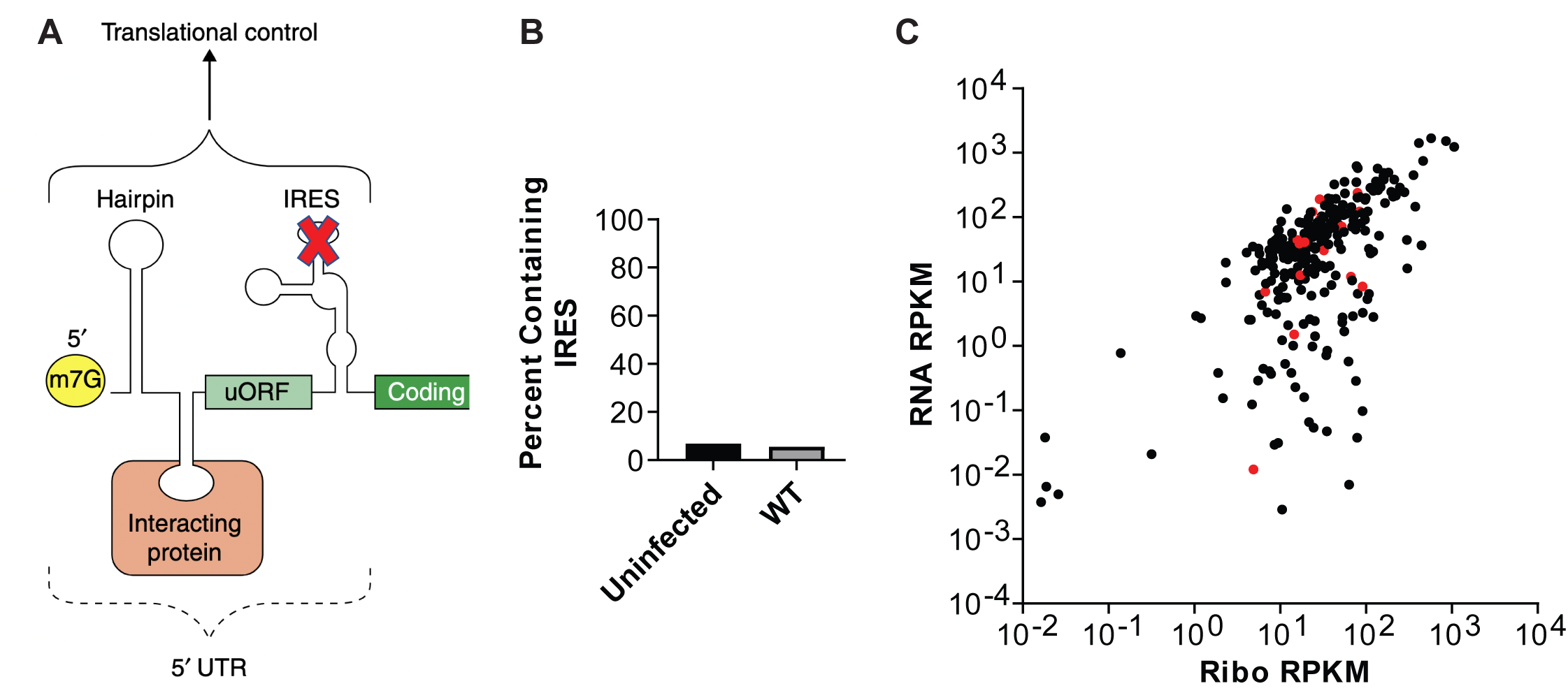
IRES are not enriched in MS candidates. (A). Cartoon showing all potential structures in 5’ untranslated region (X: transcripts missing IRES). Reprinted by permission from Springer Nature, <u>Genome Biology</u>, Untranslated Regions of mRNAs, Mignone, F., *et al*. 2002 [76]. (B) 5’ UTRs were identified having IRES sequences in MS candidates from uninfected cells and those challenged with *L. pneumophila* WT. (C). Ribosome loading of 5’ UTRs having IRES sequence in MS candidates from *L*.*pneumophila*-infected cells. Displayed are IRES-containing (Red) and IRES-absent (Black) 5’UTRs.

**Supplementary Figure 3.**
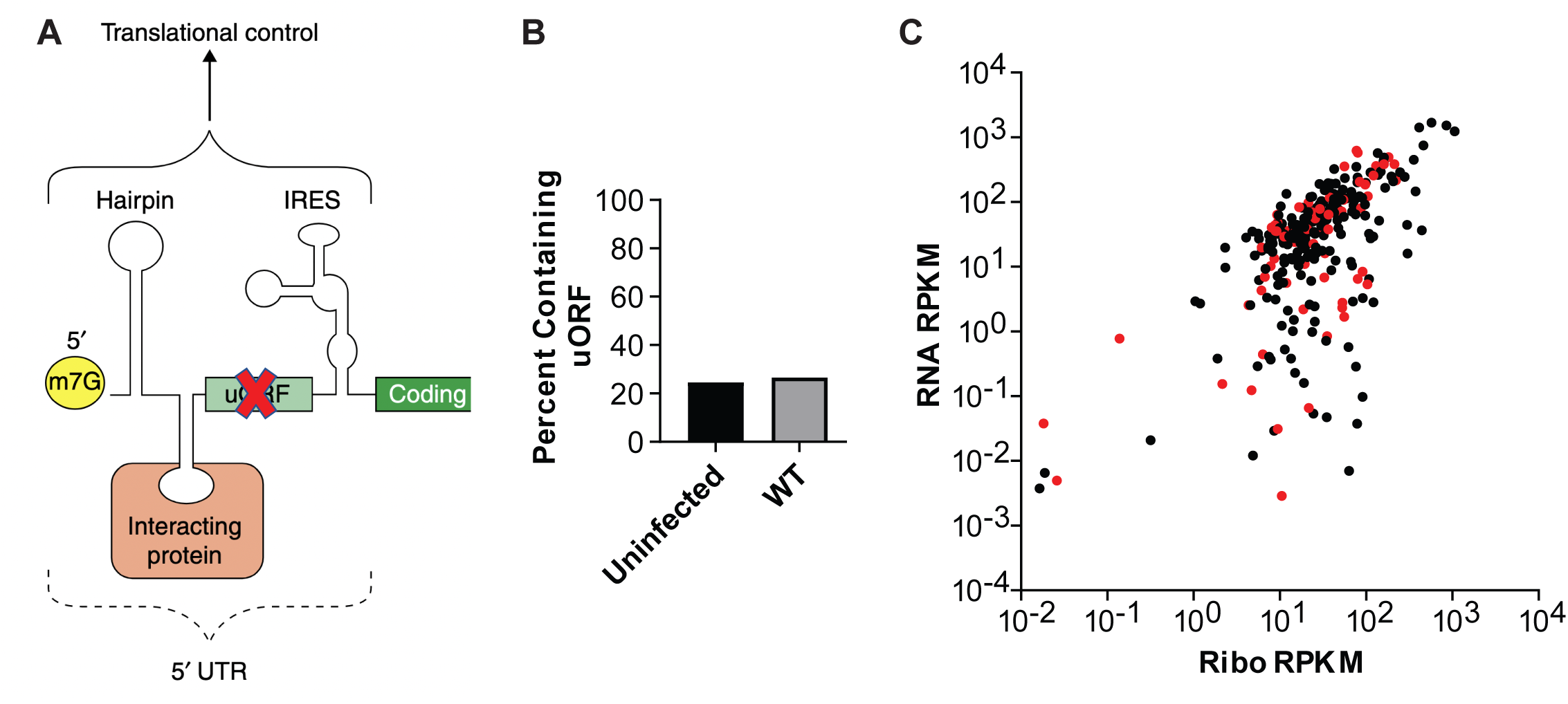
uORFs are not enriched in MS candidates. (A). Cartoon showing all potential structures in 5’ untranslated region (X: transcripts missing IRES and or uORFs) Reprinted by permission from Springer Nature, <u>Genome Biology</u>, Untranslated Regions of mRNAs, Mignone, F., *et al*. 2002 [76] (B). 5’ UTRs were identified having uORF sequences in MS candidates from uninfected cells and those challenged with *L. pneumophila* WT. (C). Ribosome loading of 5’ UTRs having uORF sequence in MS candidates from *L*.*pneumophila*-infected cells. Displayed are uORF-containing (Red) and IRES-absent (Black) 5’UTRs.

**Supplementary Table 1:**
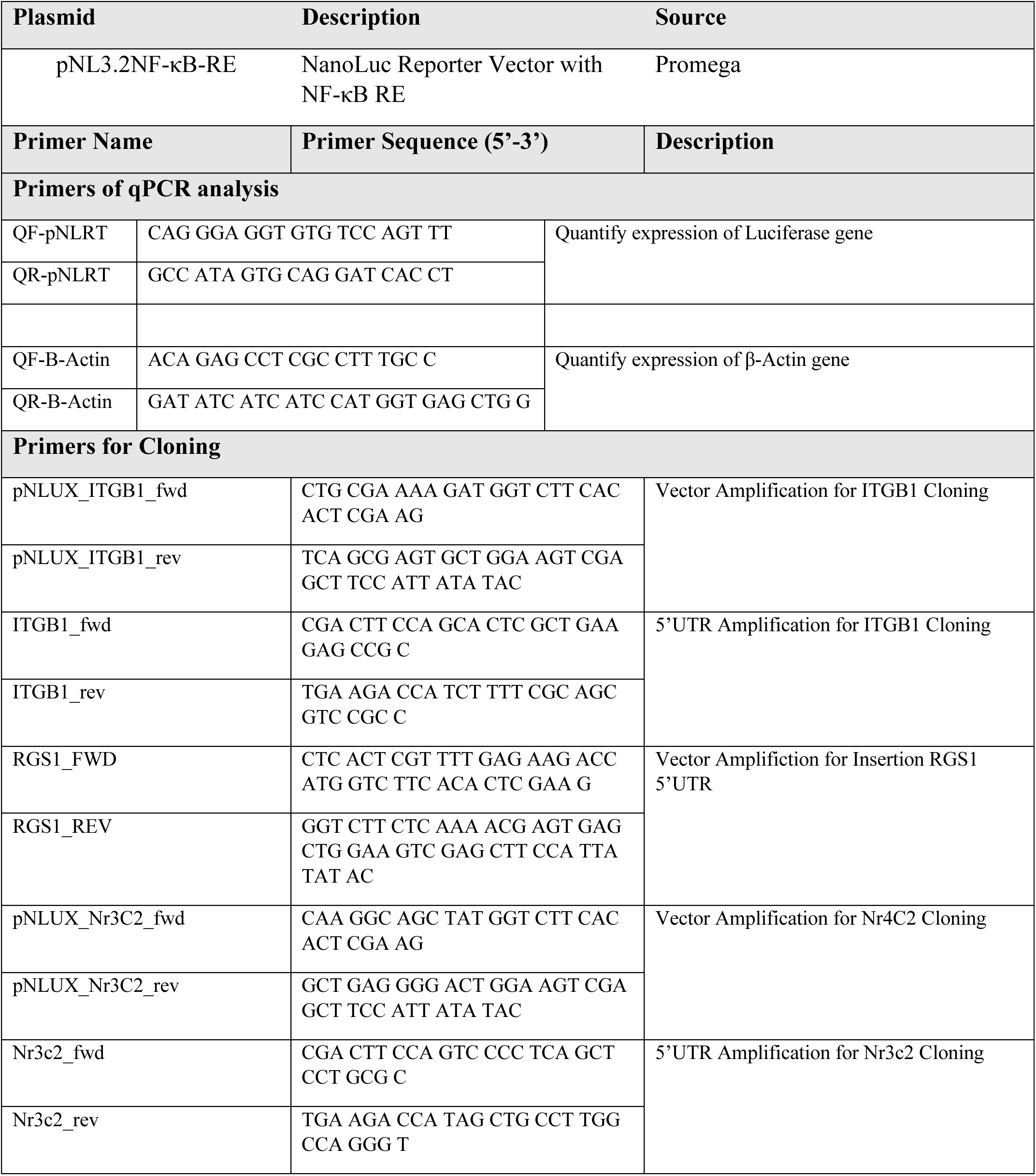
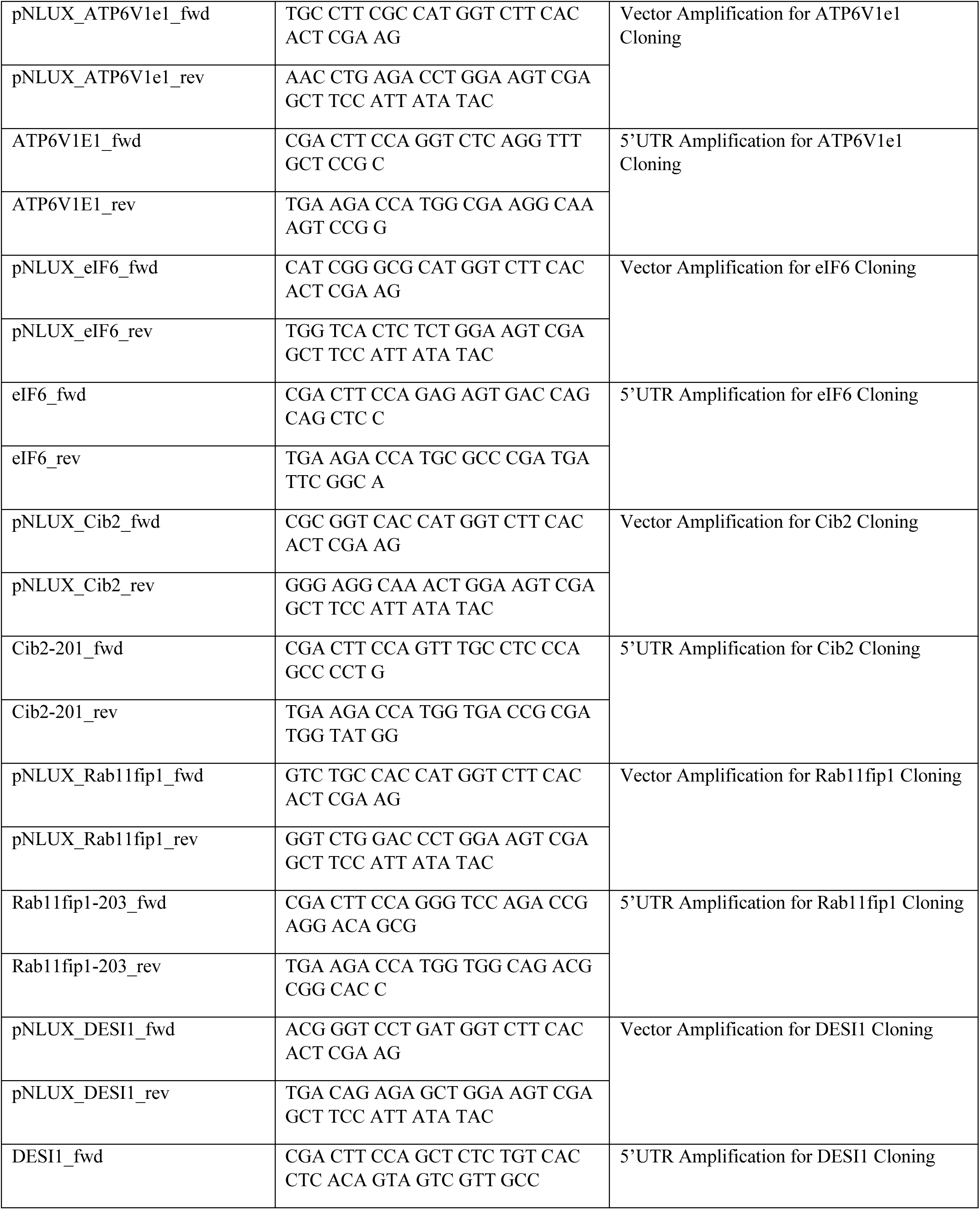
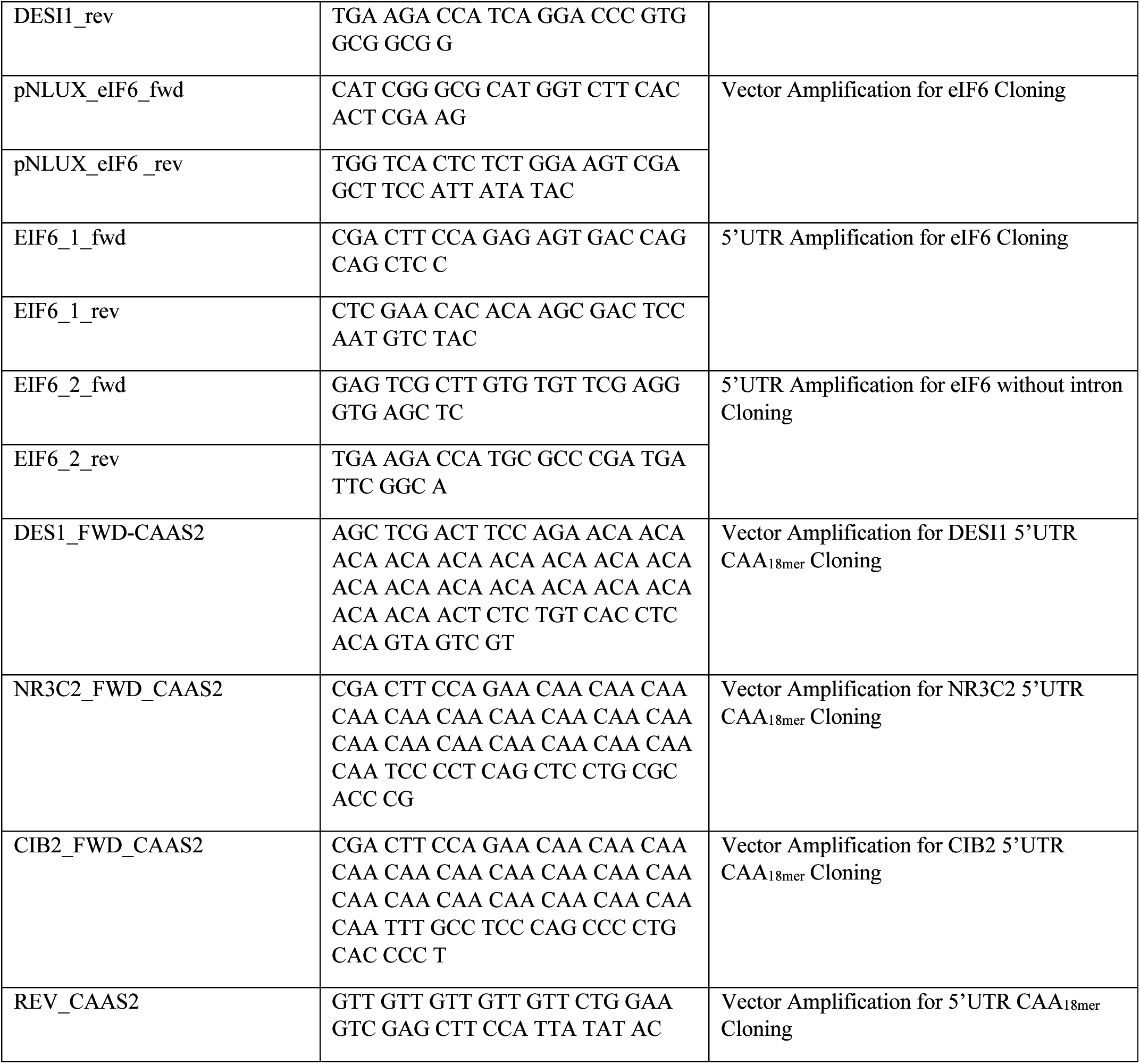
Plasmids and Primers used in this study.

